# Pacing behaviour in laboratory macaques is an unreliable indicator of acute stress

**DOI:** 10.1101/568378

**Authors:** Colline Poirier, Caitlin J. Oliver, Janire Castellano Bueno, Paul Flecknell, Melissa Bateson

## Abstract

Pacing behaviour, the most frequent stereotypic behaviour displayed by laboratory rhesus macaques (*Macaca mulatta*) is often used as an indicator of stress. In this study, we investigated how reliable this welfare indicator is at detecting acute stress by testing the reaction of macaques to the stressful event of being exposed to an agonistic interaction between conspecifics housed in the same room but in a different cage. Pacing, agitated locomotion, and stress-related displacement behaviours were quantified before, during and after agonistic interaction exposure, based on video recordings of 13 socially-housed macaques in their home cage. Displacement behaviours increased after agonistic interaction exposure, confirming that the events were experienced as stressful by the focal individuals. The occurrence of pacing did not increase during or after the agonistic interactions. Instead, agitated locomotion increased during the agonistic interactions. These results suggest either, that pacing as an indicator of acute stress is prone to false negative results, increasing in some stressful situations but not others, or that agitated locomotion has been mistaken for pacing in previous studies and that pacing is in fact unrelated to current acute stress. Both interpretations lead to the conclusion that pacing is unreliable as an indicator of acute stress in laboratory rhesus macaques.

---

Of the laboratory animal species commonly used in biomedical research, non-human primates (NHPs) are phylogenetically closest to humans. This proximity makes NHPs a crucial animal model in biomedical science, but also makes their welfare of particular concern to the general public. Current regulations reflect this public concern^1,2^, and require that researchers ensure the best possible welfare for the animals that they use. To achieve this goal, reliable welfare indicators are necessary. Stereotypies, which are repetitive, unvarying and apparently functionless behaviours, are often used as indicators of acute or chronic stress^3^. In rhesus macaques, one of the main NHP models used in research, pacing is the most common stereotypic behaviour observed in the laboratory^4^. In this study, we sought to establish whether pacing is a reliable indicator of acute stress in this species.

Pacing in rhesus macaques (hereafter macaques) is defined as repetitive walking of an individual in the exact same pattern^4,5^. Previous studies have reported that macaques increase their pacing during or immediately after short periods of acute stress (e.g. placing in a transport box for few seconds, exposure to a 10-minute cue of impending social isolation)^6,7^. Injecting macaques with a high dose of the anxiogenic drug FG7142 also increases pacing^8^. An increase in pacing frequency is thus usually interpreted as an increase in acute stress^9,10^, where the term ‘stress’ is used to refer to an unpleasant, arousing affective state.

However, in a recent study where macaques were submitted to the human intruder test, a classical test to induce acute stress, pacing frequency was found to decrease during the stressful event^11^. This result suggests that pacing can increase or decrease with acute stress, depending on the stressful situation. This finding is important from a welfare perspective, because if the frequency of a behaviour can increase in some stressful situations and decrease in others, variation in the behaviour in a new situation (whose stressful nature is unknown) cannot be interpreted unequivocally; in other words, the behaviour is unreliable as an indicator of acute stress.

In the current study we sought to determine whether the behavioural response to the human intruder test is an exceptional case, or whether other types of stressors also induce a decrease in pacing. To do this, we needed to identify an alternative source of acute stress that occurs relatively frequently in laboratory macaques and hence had the potential to explain the prevalence of pacing in primate research facilities. We chose to investigate the effect of passive exposure to agonistic interactions between conspecifics on the a priori assumption that this is stressful. Agonistic interactions in macaques are used to challenge or reassert the social position of protagonists and can result in severe injuries. In free-ranging macaque colonies, agonistic support (where one or several individuals help one of the protagonists) is frequent^12^. Hence, witnessing an agonistic interaction implies potential imminent involvement in a stressful situation, which is likely to be stressful in itself. Witnessing agonistic interactions between macaques is also a source of transient negative arousing emotion in human staff, despite years of experience with the phenomenon. Exposure to agonistic interactions also has ecological relevance, since it occurs in research colonies where individuals are housed in pairs or small social groups, which is now the norm in European primate research facilities and an emerging trend in North America. Guided by ethical considerations, we made use of agonistic interactions that occurred spontaneously in our research colony. We measured the occurrence of pacing before and during passive exposure to agonistic interactions between conspecifics housed in the same room but in a different cage than focal individuals. In the recent intruder test study^11^, the authors did not include behavioural observation of macaques after exposure to the acute stressor had ceased. It is thus possible that an increase of pacing frequency happened subsequently and was not recorded. To control for the possibility that the expected increase in pacing could be delayed, we also measured the occurrence of pacing after exposure to the agonistic interactions.

## Results

### Agonistic interactions

Video footage of home-cage behaviour of 13 focal adult male individuals previously recorded for another purpose was used for this project. Agonistic interactions were defined as any period lasting at least 10 s during which threatening/alarm vocalisations associated with loud object banging noises were heard. This identification was made based on the soundtrack of the video recordings with the video images concealed in order to avoid any selection bias caused by the behaviour of the focal individuals. Fourteen agonistic interactions were identified. The maximal duration of the interactions was 20s. Analyses of video images confirmed that in all but one case agonistic interactions occurred in a different cage than those of focal individuals. Data related to that agonistic interaction were discarded. Video recordings of each focal individual were not always available for each agonistic interaction. The number of agonistic interactions for which video existed varied between 4 and 12 agonistic interactions per individual, resulting in a total of 106 behavioural responses to agonistic interactions occurring in a cage other than that of the focal individual.

### Displacement behaviours

To verify that agonistic interaction exposure was experienced as stressful by the macaques, we recorded and analysed the occurrence of displacement behaviours^13^ displayed by macaques, namely scratching, body shaking and self-grooming (Table 1). These behaviours were chosen based on their pharmacological validation as indicators of stress or anxiety by Schino and colleagues^14^. Exposure to a stressful event induces an immediate increase in catecholamines and a slower increase in cortisol, peaking around 20 to 30 minutes after the stressful event^15–17^. To capture the behavioural responses potentially associated with these two distinct hormonal responses, behaviour of focal individuals was scored in four time intervals relative to the agonistic interaction: during the 15 min preceding the agonistic interaction (the time interval designated [−15 – 0]), during the agonistic interaction (designated as [0]), during the 15 min following the end of the agonistic interaction (designated as [0 – 15]), and during the time interval [15 – 40] min post-agonistic interaction. The four time intervals were divided into 10-s time bins and the presence (1) or absence (0) of displacement behaviours within each time bin was recorded. Following Schino and colleagues^14^ approach, scratching, body shaking and self-grooming were then combined to create a single category of ‘displacement behaviours’. Presence of at least one of the behaviours (1) or absence of all of them (0) within each time bin was computed and analysed.

**Table 1:**
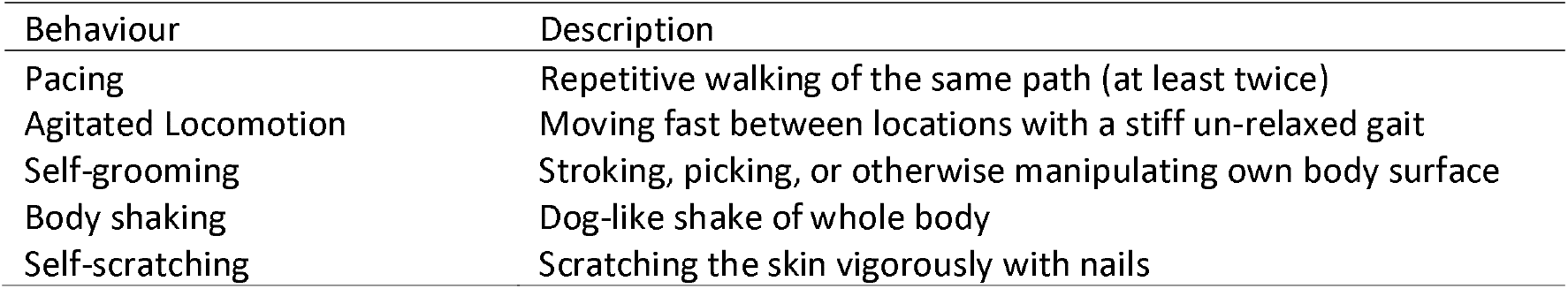
Ethogram

Displacement behaviours were displayed by all individuals, with occurrence varying between 5.5 and 27.2 % (percentage of 10-s time bins in which displacement behaviours occurred). Generalised linear mixed models of displacement data revealed an effect of time interval relative to agonistic interaction (Table 2, model 1 vs. model 2: *χ^2^* (3) = 31.1, *P* < 0.001), driven by an increase in displacement behaviour occurrence in the second time interval after the agonistic interaction, compared to before (Table 2, model 2: [15 – 40] vs. [−15 – 0]: *β* (s.e.) = 0.24 (0.05), *Z*= 4.9, *P*< 0.001).

**Table 2:**
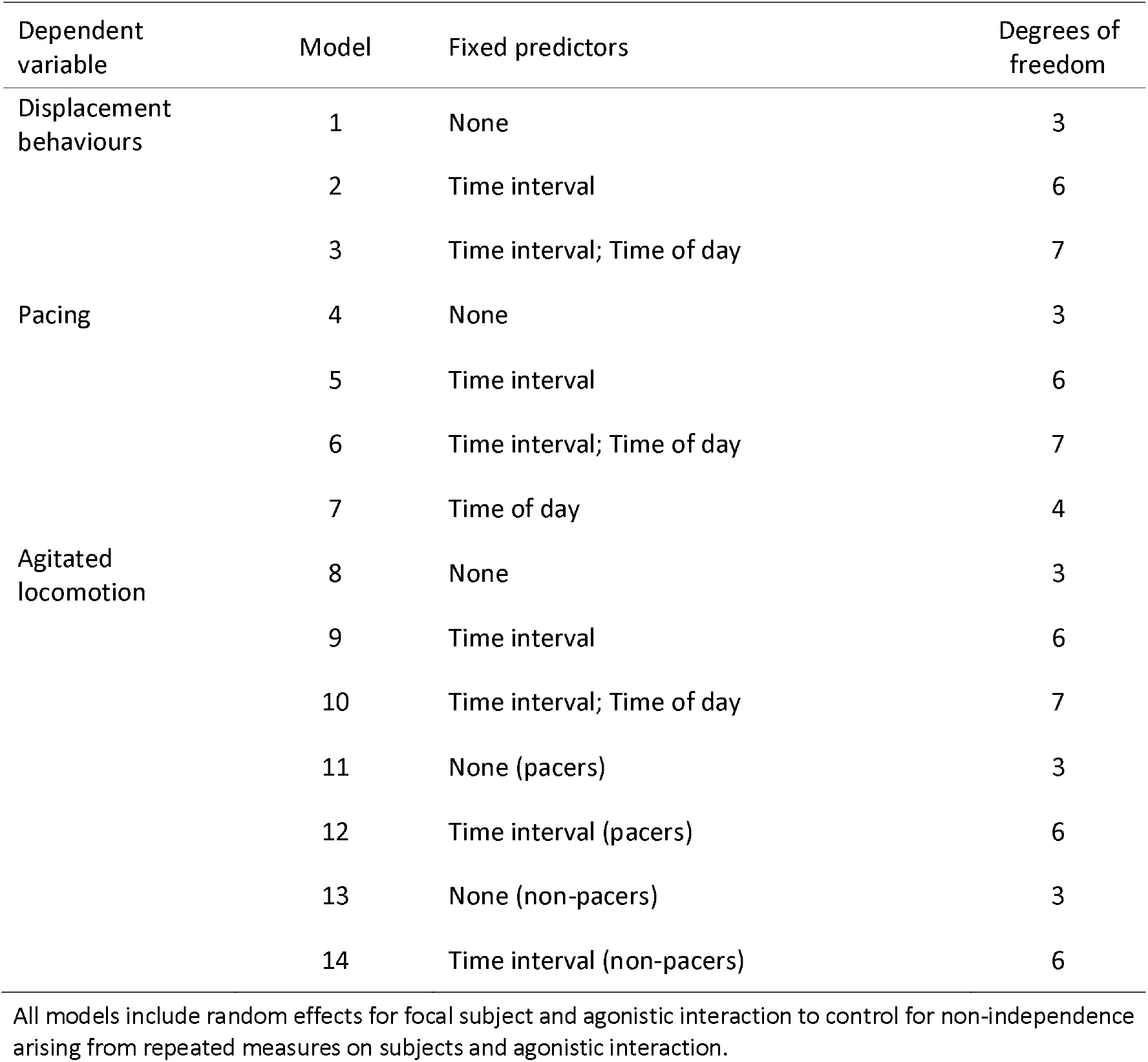
Generalised linear mixed models

| Dependent variable | Model | Fixed predictors | Degrees of freedom |
| --- | --- | --- | --- |
| Displacement behaviours | 1 | None | 3 |
|  | 2 | Time interval | 6 |
|  | 3 | Time interval; Time of day | 7 |
| Pacing | 4 | None | 3 |
|  | 5 | Time interval | 6 |
|  | 6 | Time interval; Time of day | 7 |
|  | 7 | Time of day | 4 |
| Agitated locomotion | 8 | None | 3 |
|  | 9 | Time interval | 6 |
|  | 10 | Time interval; Time of day | 7 |
|  | 11 | None (pacers) | 3 |
|  | 12 | Time interval (pacers) | 6 |
|  | 13 | None (non-pacers) | 3 |
|  | 14 | Time interval (non-pacers) | 6 |

All models include random effects for focal subject and agonistic interaction to control for non-independence arising from repeated measures on subjects and agonistic interaction.

Despite the fact that our analyses were restricted to a narrow time window of 55 min, circadian changes might still potentially confound the effect of time intervals relative to agonistic interaction. Based on unpublished data suggesting that some behaviours might follow a circadian rhythm, we thus performed a second analysis where the effect of absolute time of the day was included. Adding the time of the day as an additional fixed effect did not improve model fit (Table 2, model 2 vs. model 3: *χ^2^* (1) = 0.2, *P* = 0.67). Additional control analyses are described in the supplementary note.

### Pacing

Following the same approach as for displacement behaviours, presence or absence of pacing within each time bin was recorded (for the operational definition of pacing, see table 1). Pacing behaviour was displayed by 7 out of 13 individuals. Among pacing individuals, occurrence of pacing varied from 0.4 to 44.2 % (percentage of 10-s time bins in which pacing occurred).

Generalised linear mixed models of pacing data revealed a significant effect of time interval relative to agonistic interaction (Fig. 1; Table 2, model 4 vs. model 5, *χ^2^* (3) = 74.3, *P*< 0.001), driven by a decrease in occurrence of pacing in the two time intervals following the agonistic interactions compared to before (Table 2, model 5: [0 – 15] vs. [−15 – 0]: *β* (s.e.) = −0.24 (0.07), *Z* = −3.3, *P*< 0.001; [15 – 40] vs. [−15 – 0]: *β* (s.e.) = −0.56 (0.07), *Z* = −8.5, *P*< 0.001). There was no difference in occurrence of pacing before and during the agonistic interactions (Table 2, model 5: *β* (s.e.) =0.002 (0.34), *Z* = 0.007, *P* = 0.99).

**Figure 1:**
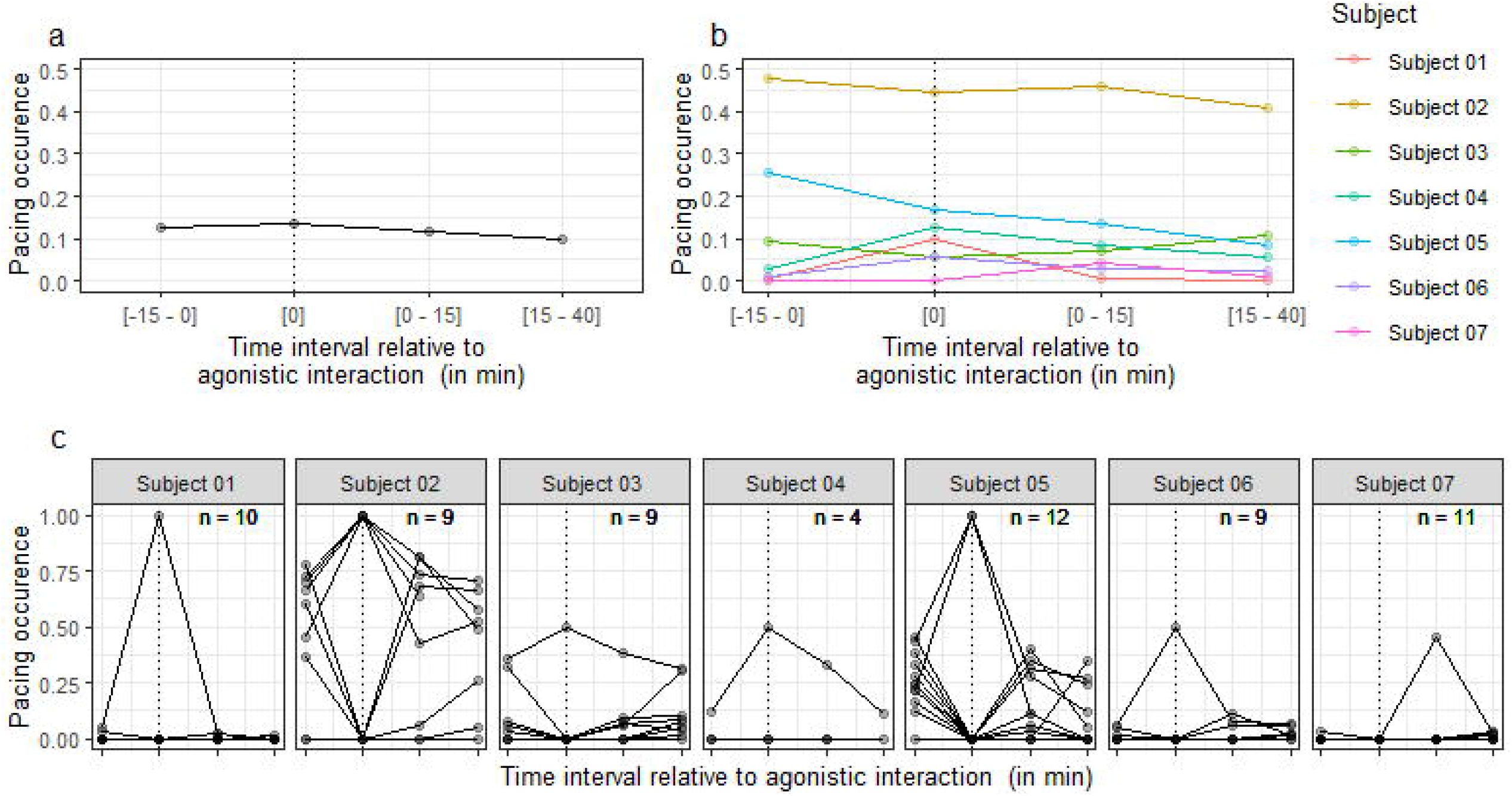
Occurrence of pacing as a function of the time interval relative to agonistic interaction. Occurrence of pacing corresponds to the proportion of 10-s time bins when the behaviour was displayed (with 1 corresponding to 100 %). The dotted line indicates the ‘during agonistic interaction’ interval. a: Data averaged over agonistic interactions and subjects. b: Data averaged over agonistic interactions. c: Data per agonistic interaction and per subject. Note that points appear darker when data from several agonistic interactions are superimposed. The n value indicated in the corner of each individual plot corresponds to the number of agonistic interactions. For the legend of the x axis, see plots a and b.

We also tested the possibility that the decrease in pacing might be due to an effect of absolute time of the day. Including time of day as an additional fixed effect significantly improved model fit (Table 2, model 5 vs. model 6, *χ^2^* (1) = 7.1, *P* = 0.007). Occurrence of pacing was found to decrease with time of day (Table 2, model 6: *β* (s.e.) = −16.63 (5.55), *Z* = −3.0, *P* = 0.002) and the effect of time interval relative to agonistic interaction in the new model was no longer significant (Table 2, model 6 vs. model 7: *χ^2^* (3) = 1.1, *P* = 0.79).

### Agitated locomotion

Finally, we also assessed the occurrence of agitated locomotion, a behaviour defined as ‘moving rapidly between locations, with a stiff un-relaxed gait’ (Table 1). In our macaque facility where individuals are housed in large and enriched cages, this behaviour differs from pacing not only in the gait of the subject (which is stiff in agitated locomotion and elastic in pacing) but also in the flexibility of the path used within and between occurrences, making the two behaviours easy to distinguish (the inter-observer reliability measured with Kappa scores were of 0.98 for pacing and 1 for agitated locomotion in the present study). Based on past cage-side observations from researchers and technicians working in the facility, we had been discussing whether this behaviour could be an indicator of stress (see ^18^ for the first study where this behaviour was systematically quantified in our facility, with inconclusive results in a context of chronic stress). In the present study, we tested the hypothesis that agitated locomotion might increase during or after exposure to an acute stressor.

Agitated locomotion was displayed by 12 out of 13 individuals, including all pacers (*N* = 7) and 5 non-pacers. The occurrence of agitated locomotion among individuals that displayed the behaviour at least once varied from 0.03 to 1.3 % (percentage of 10-s time bins in which agitated locomotion occurred).

Generalised linear mixed models of agitated locomotion data revealed a significant effect of time interval relative to agonistic interaction (Table 2, model 8 vs. model 9: *χ^2^* (3) = 154, *P*< 0.001), driven by an increase in agitated locomotion during agonistic interactions, compared to before (Table 2, model 9: [0] vs. [−15 – 0]: *β* (s.e.) = 4.1 (0.29), *Z*= 14.4, *P*< 0.001). There was no significant difference in occurrence of agitated locomotion before and after the agonistic interactions (Table 2, model 9: [0 – 15] vs. [−15 – 0]: *β* (s.e.) = −0.56 (0.30), *Z* = −1.90, *P* = 0.06; [15 – 40] vs. [−15 – 0]: *β* (s.e.) = 0.003 (0.23), *Z* = 0.01, *P* = 0.99). Adding the absolute time of day as an additional fixed effect did not improve model fit (Table 2, model 9 vs. model 10: *χ* (1) = 0.27, *P* = 0.60).

Generalised linear mixed models applied separately to pacers and non-pacers revealed similar effects of time interval relative to agonistic interaction in both groups (Fig. 2 and Table 2, pacers: model 11 vs. model 12: *χ^2^* (3) = 124, *P*< 0.001, model 12: [0] vs. [−15 – 0]: *β* (s.e.) = 4.39 (0.34), *Z*= 12.7, *P*< 0.001; non-pacers: model 13 vs. Model 14: *χ* (3) = 33, *P*< 0.001, model 14: [0] vs. [−15 – 0]: *β* (s.e.) = 3.49 (0.59), *Z*= 5.9, *P*< 0.001).

**Figure 2:**
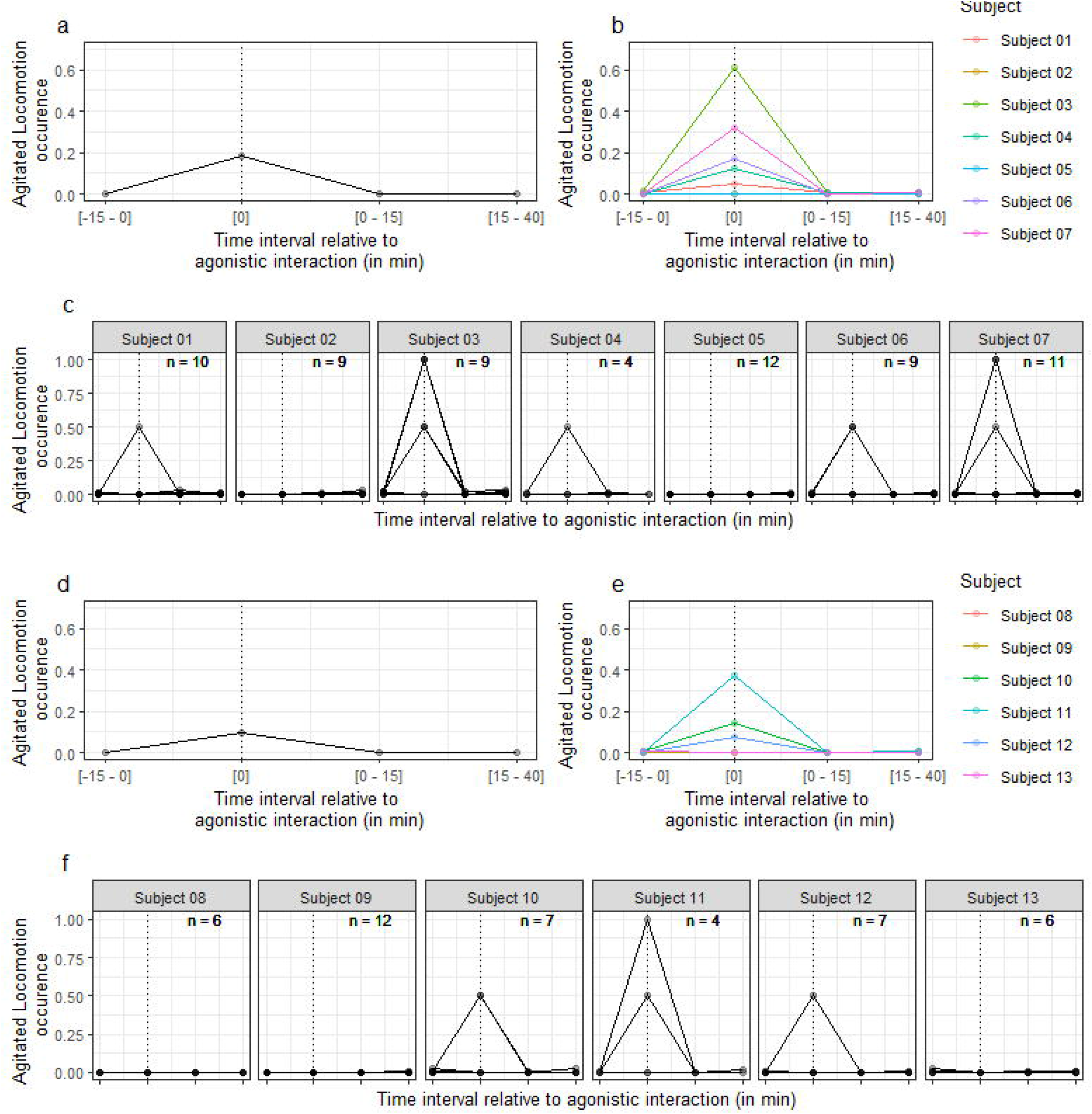
Occurrence of agitated locomotion as a function of the time interval relative to agonistic interaction in pacers (a-c) and non-pacers (d-f). Occurrence of agitated locomotion corresponds to the proportion of time bins when the behaviour was displayed (with 1 corresponding to 100 %). The dotted line indicates the ‘during agonistic interaction’ interval. a/d: Data averaged over agonistic interactions and subjects. b/e: Data averaged over agonistic interactions. c/f: Data per agonistic interaction and per subject. Note that points appear darker when data from several agonistic interactions are superimposed. The n value indicated on each plot corresponds to the number of agonistic interactions. For the legend of the x axis, see plots a/d and b/e.

## Discussion

This study investigated the behavioural reaction of laboratory rhesus macaques to agonistic interactions occurring between conspecifics housed in separate, but nearby cages within the same room. Displacement behaviours, which are pharmacologically validated indicators of stress or anxiety, increased after the agonistic interactions, confirming that witnessing agonistic interactions was perceived as stressful by the macaques. Our study reveals that stereotypic pacing did not increase during or after the agonistic interactions. Peterson and colleagues previously showed that pacing did not increase during an intruder test, a paradigm established to induce stress in macaques^11^. Here we extend this result by testing a different stressful situation (passive exposure to an agonistic interaction between conspecifics) and by providing a more comprehensive assessment of pacing by testing its frequency not only during the stressful event but also during a sustained period of time (up to 40 min) after the stressful event. These data contrast with previous studies where pacing was found to increase with acute stress^6,7^. These conflicting results can be interpreted in two different ways that we describe in the following paragraphs.

The first possible explanation for discrepant results from different studies is that pacing frequency increases in some stressful situations but not in others. If this is the case, it means that pacing when used as an indicator of acute stress is prone to false negative results, where macaques experience acute stress but do not express it by an increase in pacing (Fig. 3). From a welfare perspective, this possibility is problematic for two reasons. First, using pacing as an indicator of acute stress might lead stressful husbandry or experimental procedures to fail to be identified as such, preventing researchers, veterinarians and/or technical staff from refining them and thus improving animal welfare. Second, false-negative results also undermine the use of an absence of pacing frequency increase during or after a procedure as a reliable indicator that the procedure is not stressful. This is problematic because determining which husbandry and scientific procedures are not experienced as stressful is as important as identifying those which are stressful. It should also be noted that whether pacing in macaques increases after positively-valenced (i.e. pleasant) arousing events has never been investigated. The potential of pacing to generate false positive results (Fig. 3) when used as an indicator of acute stress is thus currently unknown. Consequently, when pacing frequency is found to be increased by an event or procedure, one cannot conclude with confidence that this event or procedure induced acute stress rather than a pleasant arousing emotion.

**Figure 3:** Different possible relationships between pacing frequency and acute stress.

Alternatively, the fact that pacing seems to increase during some stressful situations^6,7^, but not others (^11^, our study), might be due to the possible conflation between pacing and agitated locomotion in previous studies. Indeed, our study revealed an increase in agitated locomotion during agonistic interactions. Differences between pacing and agitated locomotion are summarised in Table 3. One notable difference is that agitated locomotion is not stereotypic, the path used by an individual varying from one occurrence to another. In our research facility where macaques are housed in relatively big cages (15 m^3^) with shelves and ropes, the two behaviours are easy to distinguish. However, it is possible that in small cages agitated locomotion appears stereotypic due to the lack of options in the paths individuals can walk, making it visually indistinguishable from true pacing. Interestingly, studies reporting an increase in pacing frequency induced by acute stress, including the anxiogenic drug FG7142^6–8^, all come from macaques housed in very small cages (below 1.3 m^3^). It is thus possible that existing literature linking acute stress and pacing should be reinterpreted as showing a link between acute stress and agitated locomotion. This hypothesis raises the possibility that pacing displayed by macaques in relatively big cages is unrelated to present acute stress.

**Table 3:** Differences between Pacing and Agitated Locomotion

|  | **Pacing** | **Agitated locomotion** |
| --- | --- | --- |
| *Form* | Elastic gait | Inelastic gait |
| *Flexibility* | Invariant path | Variable path |
| *Prevalence*<sup>1</sup> | About half of individuals (7/13) | Almost all individuals (12/13) |
| *Frequency*<sup>1</sup> | Frequent (up to 44.2%) | Rare (up to 1.3 %) |
| *Duration*<sup>1</sup> | Long (up to 10 min) | Short ($\leq 20$ s) |
| *Context*<sup>1</sup> | Before, during and after agonistic interactions | Almost exclusively during agonistic interactions |

<sup>1</sup> in this study

In some individuals, occurrence of pacing was found to be high before, during and after the agonistic interactions (fig. 1B). Source(s) of acute stress during the whole period could not be identified. This result indicates that pacing is likely to be caused by factor(s) other than current acute stress. We have recently reviewed these potential factors^19^. Our review suggests that pacing might be the consequence of brain abnormalities induced by a past stressful event, and be unrelated to the current state of the individual. Alternatively, pacing might be caused by chronic stress, boredom or a specific need to walk. These different hypotheses have different welfare consequences with the welfare of pacing macaques being either better, worse or equivalent to that of non-pacing individuals^19^. As long as the cause(s) of pacing remain unresolved, we argue that pacing should not be used as an indicator of acute or chronic stress, especially in the absence of other indicators of poor welfare.

Our results raise the possibility that agitated locomotion could be a useful indicator of acute stress. In sharp contrast with pacing behaviour, agitated locomotion increased during the agonistic interactions but was virtually non-existent outside periods of agonistic interactions, when no source of acute stress could be detected. These results however need to be replicated with other stressful situations. If agitated locomotion is systematically observed during different stressful situations (i.e. if no/few false negative results are observed), this behaviour might be a convenient welfare marker since it is easy to identify (at least in large, enriched cages) and its baseline frequency near to zero makes any increase easy to detect. Whether agitated locomotion also increases during positively-valenced arousing events (susceptibility to false positive results) will also need to be determined.

To our knowledge, the precise time scale of behavioural indicators of acute stress in macaques has never been systematically investigated. From an applied perspective, this information might help in designing better behavioural observation protocols. At the physiological level, acute stress induces a well-established cascade of events, including the release of different hormones over different time scales. In the present study, we used this knowledge to guide our observation periods. Agitated locomotion increased during the very brief period of agonistic interactions, suggesting that this behaviour might be triggered by the fast-released catecholamines. By contrast, displacement behaviours’ increase was observed during the 15 - 40 minute interval after the stressor, suggesting that these behaviours might be induced by the slowly-released cortisol. We did not quantify the occurrence of behaviours later than 40 minutes post-stressor, and therefore, cannot exclude that pacing might potentially increase at this much later time point. Further studies will be necessary to determine whether quantifying pacing several hours after a stressful event could be useful as an indicator of acute stress.

To conclude, by showing that pacing does not increase following exposure to conspecific agonistic interactions, this study shows that pacing behaviour is not a reliable indicator of acute stress. Our data also suggest that agitated locomotion might potentially be a useful indicator of acute stress, or at least arousal, but more studies are needed to confirm this. Welfare research on captive animals is often done using experimental paradigms that involve manipulating the affective state of the animals by imposing a stressor^20,21^. While this approach is undoubtedly the most powerful, the current study illustrates how new knowledge can also be gained by using observations of animals without inducing additional stress.

## Materials and methods

### Subjects and ethical statement

Subjects were 13 male rhesus macaques (*Macaca mulatta*) aged between 4 and 10 years (weight range: 5 to 18 kg). Individuals had been raised in a British breeding centre, first with their mother for at least 6 months and then with other juveniles in large rooms. When adolescent, subjects were moved to the Newcastle University research facility, which complies with the NC3Rs Guidelines for ‘Primate Accommodation, care and use’ (www.nc3rs.org.uk/primatesguidelines). There, they were housed in relatively large cages (2.1 × 3.0 × 2.4 m) exceeding the minimal space requirement under the UK legislation (1.8 m^3^ per animal). The height of the cages and the presence of high perches and shelves allowed macaques, if alarmed, to flee upwards. Subjects were housed in iso-sex pairs except one male, which was paired with two females. Besides the presence of at least one cage mate, enrichment was provided by daily foraging opportunity (the food was scattered in the litter on the floor), wooden shelves, swings, ropes, objects to manipulate (changed on a regular basis) and natural light. Cages were located in a large room housing more than 40 individuals. Focal individuals were thus in visual and auditory contact with many other conspecifics in addition to their cage mates. Some of the individuals were routinely involved in neuroscience experiments while others had not yet been subjected to any experimental procedures. This study consisted of behavioural observation of macaque home cage behaviour recorded via remotely-controlled cameras, inducing no interference with the animals. The stressor consisted of passive exposure to agonistic interactions occurring naturally between other animals; the frequency of agonistic interactions was not manipulated for this study. No licence was required for this study.

### Behavioural observation

Videos were recorded with remotely-controlled cameras (Cube HD 1080, Y-cam Solutions Limited, Twickenham, UK). Recordings from between 1600 and 1800 on Saturdays and Sundays between January and December 2015 were analysed. This period of the week was chosen as it was the only period when macaques were in their home cage with their cagemate, awake and no people were present in the facility. This strategy avoided the necessity of having to control for the potentially stressful effects of human presence and cage mate absence, which typically happens on a daily basis during week days when subjects are involved in experiments.

### Behavioural scoring and analysis

Behavioural scoring was performed by two observers, including one blind to the hypotheses under test and to when the agonistic interactions occurred. Measures of inter-rater reliability revealed kappa scores above 0.96. Statistical analyses were performed in R, version 3.3.2^22^ using the *lme4* package^23^. The presence or absence of each behaviour within each 10-second time bin was analysed using generalised linear mixed models (*glmer* function) with binomial error distribution and logistic link function. The *time interval relative to agonistic interaction* was declared as a fixed effect (categorical effect with 4 levels), while *subjects* and *agonistic interactions* were declared as random effects, with *agonistic interactions* being nested within *subjects*. Maximum-likelihood estimation was employed throughout. Note that these models cope with highly unbalanced designs such as this one (the amount of data in the [0] interval being much less than in the other time intervals) and non-independence due to repeated measures of subjects and agonistic interactions. Significance testing was carried out by the likelihood ratio test, which compares the change in deviance when a term is excluded from the model with the χ^2^ distribution.

### Data availability

The datasets generated during and/or analysed during the current study are available from the corresponding author on reasonable request.

## Supporting information

Supplementary results

## Acknowledgements

This work was supported by the Association for the Study of Animal Behaviour, the NC3Rs (NC/K000802/1) and internal funding from Newcastle University. The funders had no role in the writing of the report nor in the decision to submit the article for publication.

## Author contributions statement

Funding acquisition: C.P., P.F., M.B; Conceptualisation and Methodology: C.P and M.B.; Resources: P.F.; Investigation: C.P., J.C.B. and C.O.J.; Formal analysis: C.P.; Supervision: C.P. and M.B.; Visualisation: C.P.; Writing – original draft: C. P.; Writing – review and editing: C.P., C.O.J, P.F., M.B.

