## Supplementary results for "Pacing behaviour in laboratory macaques is an unreliable indicator of acute stress"

Age effect

We tested the hypothesis that variation in displacement behaviours could depend of the age of the subjects (range: 4 -10 years). Indeed, our sample included sexually mature males but some of them had not fully developed their secondary sexual characteristics (e.g. fully erupted and sharpened canines, broad muscular shoulders). Based on the assumption that the emotional response to agonistic interactions might differ between sub-adults (age < 6 years) and fully adult subjects (age > 6 years), we tested the effect of time intervals on displacement occurrence, adding an interaction term between age, defined as a categorical variable with two levels (below and over 6 years), and time intervals. This analysis revealed a significant interaction term (Supplementary Table S1, model 1 vs. model 2: *χ^2^* (4) = 83.5, *P* < 0.001). Post-hoc analyses revealed an effect of time interval in both sub-groups (Supplementary Table S1, sub-adults: model 3 vs. model 4: *χ^2^* (3) = 39.0, *P* < 0.001; adults: model 5 vs. model 6: *χ^2^* (3) = 74.8, *P* < 0.001). However, the effect in sub-adults was driven by an increase in displacement behaviour occurrence in the first time interval after the agonistic interaction, compared to before (Supplementary Table S1, model 4: [0 – 15] vs. [-15 – 0]: *β* (s.e.) = 0.42 (0.1), *Z* = 4.0, *P* < 0.001), while the effect in adults was driven by an increase in displacement behaviour occurrence in the second time interval after the agonistic interaction, compared to before (Supplementary Table 2, model 6: [15 – 40] vs. [-15 – 0]: *β* (s.e.) = 0.32 (0.05), *Z* = 5.9, *P* < 0.001). This analysis indicates that witnessing agonistic interactions was a source of stress for both adults and sub-adults, with the timing of the behavioural manifestation of stress differing between sub-groups.

Cage effect

Subjects were housed in groups of 2-3 individuals. In case that the behaviour of one subject influenced the behaviour of its cagemate, we performed additional analyses including ‘cage’ as an additional random effect (with ‘subjects’ being nested within ‘cage’). Including or not cage did not change the results: the effect of time interval on displacement behaviour was still driven by an increase in displacement behaviour occurrence in the second time interval after the agonistic interaction, compared to before (Supplementary Table S1, model 7 vs. 8: *χ^2^* (3) = 31.0, *P* < 0.001, model 8: [15 – 40] vs. [-15 – 0]: *β* (s.e.) = 0.24 (0.05), *Z* = 4.9, *P* < 0.001); the effect of time interval on pacing was still driven by a decrease of pacing in the two time intervals following the agonistic intervention (Supplementary Table S1, model 9 vs. 10: , *χ^2^* (3) = 74.3, *P* < 0.001, model 10: [0 – 15] vs. [-15 – 0]: *β* (s.e.) = -0.24 (0.07), *Z*  = -3.3, *P* < 0.001; [15 – 40] vs. [-15 – 0]: *β* (s.e.) = -0.56 (0.07), *Z* = -8.5, *P* < 0.001) and the effect of agitated locomotion was still driven by an increase during the agonistic interaction (Supplementary Table S1, model 11 vs. model 12: *χ^2^* (3) = 154, *P* < 0.001; model 12: [0] vs. [-15 – 0]: *β* (s.e.) = 4.1 (0.29), *Z* = 14.4, *P* < 0.001), compared to before.

Supplementary Table S1: Additional generalised linear mixed models of behaviour presence or absence within each time bin (binomial error distribution)

| Dependent variable | Model | Fixed predictors | Random effect | Degrees of freedom |
| --- | --- | --- | --- | --- |
| Displacement | 1 | Time interval | Subject/Fight | 6 |
|  | 2 | Time interval ; Time interval X Age | Subject/Fight | 10 |
|  | 3 | None (sub-adults) | Subject/Fight | 3 |
|  | 4 | Time interval (sub-adults) | Subject/Fight | 6 |
|  | 5 | None (adults) | Subject/Fight | 3 |
|  | 6 | Time interval (adults) | Subject/Fight | 6 |
|  | 7 | None | Cage/Subject/Fight | 4 |
|  | 8 | Time interval | Cage/Subject/Fight | 7 |
| Pacing | 9 | None | Cage/Subject/Fight | 4 |
|  | 10 | Time interval | Cage/Subject/Fight | 7 |
| Agitated locomotion | 11 | None | Cage/Subject/Fight | 4 |
|  | 12 | Time interval | Cage/Subject/Fight | 7 |

All models include random effects for focal subject and agonistic interaction to control for non-independence arising from repeated measures on subjects and agonistic interaction.
